# Comprehensive gene expression analysis of organoid-derived healthy human colonic epithelium and cancer cell line by stimulated with live probiotic bacteria

**DOI:** 10.1101/2024.05.23.595631

**Authors:** Akira Sen, Atsuki Imai, Kota Yanagisawa, Eiji Miyauchi, Tsukasa Oda, Fuki Sasaki, Shintaro Uchida, Takuhisa Okada, Takehiko Yokobori, Hiroshi Saeki, Toshitaka Odamaki, Nobuo Sasaki

**Affiliations:** The Laboratory for Mucosal Ecosystem Design, Institute for Molecular and Cellular Regulation, Gunma University, Maebashi, Gunma 371-8512, Japan; Next Generation Science Institute, Morinaga Milk Industry Co Ltd, Zama, Kanagawa 228-8583, Japan; Division of Gastroenterological Surgery, Department of General Surgical Science, Gunma University, Graduate School of Medicine, Maebashi, Gunma 371-8512, Japan; Initiative for Advanced Research, Gunma University, Maebashi, Gunma 371-8512, Japan

**Keywords:** Colon cancer cells, Colonic epithelium, gene expression, gut microbiome, probiotic bacteria, transcriptome profiling

## Abstract

The large intestine has a dense milieu of indigenous bacteria, generating a complex ecosystem with crosstalk between individual bacteria and host cells. *In vitro* host cell modeling and bacterial interactions at the anaerobic interphase have elucidated the crosstalk molecular basis. Although classical cell lines derived from patients with colorectal cancer including Caco-2 cells are used, whether they adequately mimic normal colonic epithelial physiology is unclear. To address this, we performed transcriptome profiling of Caco-2 and Monolayer cells derived from healthy Human Colonic Organoid (MHCO) cultured hemi-anaerobically. Coculture with the anaerobic gut bacteria, *Bifidobacterium longum* subsp. *longum* differentiated the probiotic effects of test cells from those of physiologically normal intestinal and colorectal cancer cells. We cataloged non- or overlapping gene signatures where gene profiles of Caco-2 cells represented absorptive cells in the small intestinal epithelium, and MHCO cells showed complete colonic epithelium signature, including stem/progenitor, goblet, and enteroendocrine cells colonocytes. Characteristic gene expression changes related to lipid metabolism, inflammation, and cell-cell adhesion were observed in cocultured live *Bifidobacterium longum* and Caco-2 or MHCO cells. *B. longum*-stimulated MHCO cells exhibited barrier-enhancing characteristics, as demonstrated in clinical trials. Our data represent a valuable resource for understanding gut microbe and host cell communication.

## Introduction

The large intestine, also known as the colon, is a crucial component of the human digestive system with a wide range of physiological functions vital for maintaining overall health[1]. One of the primary physiological functions of the colon is the absorption of water and electrolytes from residual undigested food materials entering the colon from the small intestine[2]. The content of the partially digested chyme that passes through the colon is concentrated through water absorption, an essential process that maintains body fluid balance and prevents dehydration.

Additionally, the colon hosts a diverse community of beneficial bacteria, known as the gut microbiota, which contribute significantly to human health[3]. These microorganisms aid in fermenting undigested carbohydrates and breaking down certain substances to produce metabolites, gasses, and vitamins[4]. The gut microbiota plays a crucial role in bolstering the immune system, protecting against harmful pathogens and maintaining the intestinal lining integrity[5]. The role of microorganisms in maintaining homeostasis and influencing various aspects of human physiology has spurred research into probiotics development.

Probiotics are live microorganisms that confer health benefits when consumed in adequate quantities, and they have emerged as a promising strategy for improving human health and well-being[6]. To promote research and development of probiotics, the physiological functions of microorganisms and their effects on the host need to be investigated [7]. However, details of the fundamental molecular mechanisms underlying host cell-bacteria interactions remain largely unknown. Recent improvements in next-generation sequencing and mass spectrometry technologies have increased interest in the trans-omics analysis of gut microbiota. However, the results only provide a correlation between changes in microorganisms and their traits and do not allow direct causal inference. Therefore, *in vitro* models are required to elucidate the detailed mechanisms of microbe-host cell interactions. However, existing models do not assess the response of normal human intestinal epithelial cells because of the difference in oxygen demands between gut bacteria (most bacteria living in the colon are anaerobes) and host cells (propagate under normoxia).

To overcome this shortcoming, researchers including our group, have developed a state-of-the-art coculture system of human colonic epithelium with highly oxygen-sensitive gut bacteria[8–12]. Colonic epithelial cells were cultured using a two-dimensional (2D) design on the Boyden chamber to establish the coculture system. Immortalized cell lines derived from human colorectal cancer tissue, such as Caco-2 cells, are commonly used to analyze gut bacteria and their metabolites[13]. Although the molecular basis is still unclear, the morphological and biochemical properties of Caco-2 cells are known to spontaneously transform, converting them into a small intestine-like epithelium following prolonged culture under confluent 2D layer conditions[14]. Consequently, Caco-2 cell monolayers have been extensively used as *in vitro* models of the small intestine to characterize certain aspects of nutrient uptake [15].

Most probiotic bacteria, such as *Akkermansia* and *Bifidobacterium* species, reside and function in the colon rather than the small intestine[16]. Thus, colonic epithelium must be used in analyzing the role of microbes inhabiting the colon. Recently, normal colonic epithelial cells were stably cultured for extended periods using three-dimensional (3D) mouse and human organoid culture methods[17, 18]. Self-renewing 2D <u>M</u>onolayer cultured colonic epithelium cells derived from healthy <u>H</u>uman <u>C</u>olonic <u>O</u>rganoids (MHCO) are also a physiologically relevant model[10, 19]. Moreover, primary cultured intestinal epithelial cells derived from organoids are increasingly used to study intestinal microbes.

In this study, we investigated the feasibility of using organoid-based normal human epithelium instead of Caco-2 cells in an intestinal hemi-anaerobic coculture system (iHACS)[10] containing the probiotic bacteria, *Bifidobacterium longum*. This system was established to address the limitation of currently available *in vitro* coculture systems of representative human and microbial cells. We conducted a detailed molecular analysis of the differential effects on cell physiology induced by coculture experiments with Caco-2 cells and normal colonic epithelium. By stimulating a colorectal cancer-derived cell line or normal colonic epithelium with bacteria, we aimed to generate an expression profile for the stratification of probiotic responses.

## Results

### Characterization of Caco-2 and colonic organoid-derived monolayer

We performed RNA-sequencing (RNA-seq) analysis to decipher the complete genetic information of the intestinal epithelium cultured using iHACS and compared two *in vitro* cell culture models of Caco-2 and 2D cultured normal colonic epithelium derived from MHCO. To collect RNA samples for transcriptome analysis, we first aligned the culture conditions of the Caco-2 and MHCO cells (Fig. 1A). To collect RNA samples for transcriptome analysis, we first align the culture conditions of Caco-2 and MHCO cells (Fig. 1A). Caco-2 was found to be able to grow in the organoid-conditioned medium, which is devoid of serum completely, so after 17 days of culture under sufficiently confluent conditions in fetal bovine serum-based DMEM Cell Culture Medium[20, 21]. The medium was changed to the organoid-conditioned medium, and RNA was recovered from cells for a further 3 days culture (Fig. 1A). Contrastingly, MHCO cells were cultured for 3 days after reaching confluence under similar conditions to those used for the Caco-2 cells (Fig. 1A). Importantly, we comparatively analyzed the gene expression in MHCO and Caco-2 cells following growth under hemi-anaerobic conditions.

**Figure 1.**
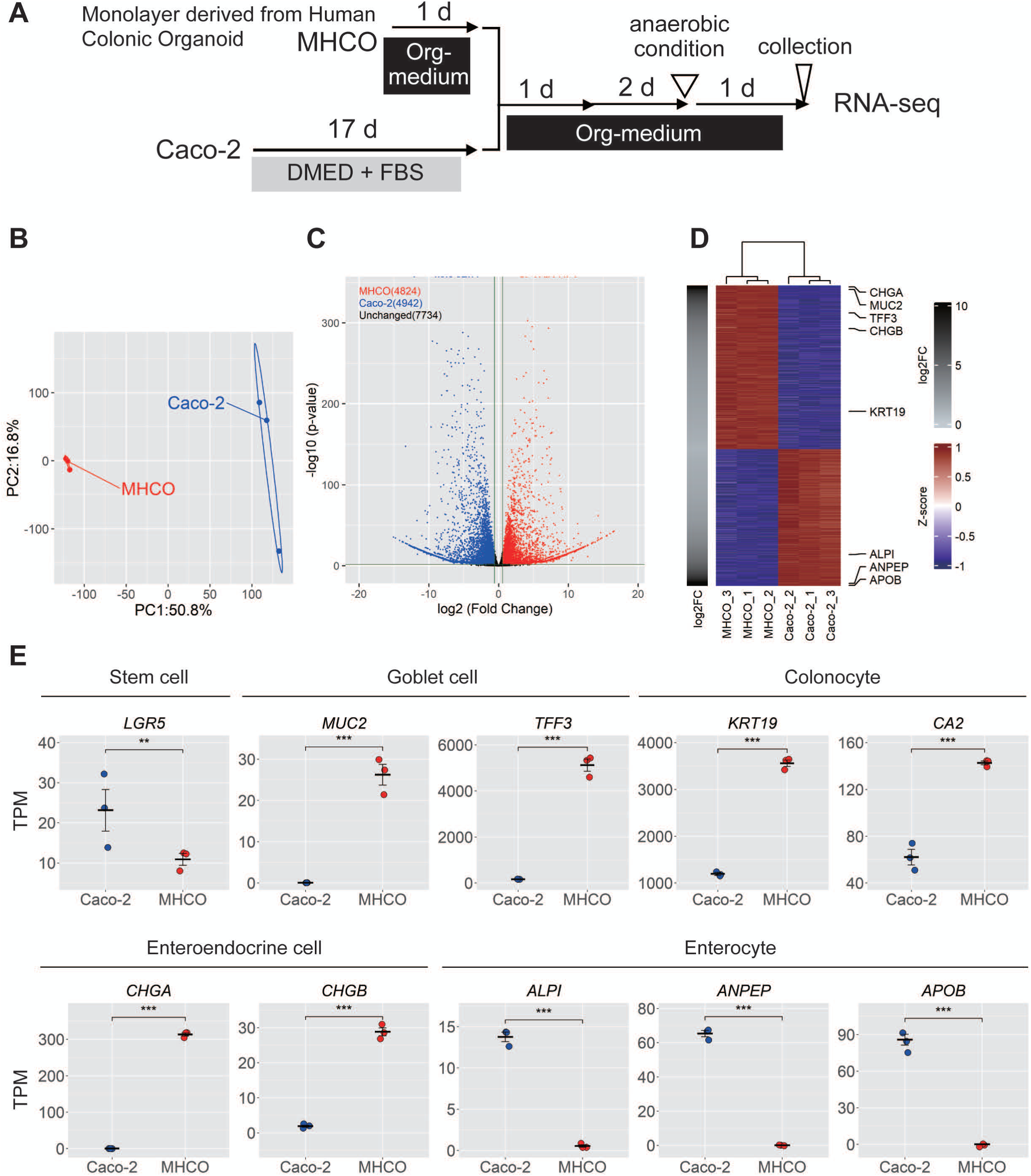
Comparison of gene expression profiles between human colonic organoid-derived monolayers (MHCO) and Caco-2. **(A)** A schematic representation of the experiment. **(B)** PCA of the transcriptomes of MHCO and Caco-2. Each plot represents a single transcriptome. Transcriptomes from three replicates were plotted. Ellipses indicate the 95% confidence interval. **(C)** Volcano plot depicting the DEGs of the transcriptomes of MHCO and Caco-2. The x- and the y-axis show the log_2_ transformed fold change and the -log_10_ transformed adjusted p-value. Red and blue dots indicate significantly upregulated genes in MHCO and Caco-2 according to adjusted p-value <0.05 and log_2_ fold change >1.5. **(D)** Heatmap depicting the top-2000 DEGs of the transcriptomes of MHCO and Caco-2 when sorted by adjusted p-value. **(E)** TPM values of marker genes for each gut epithelial cell type. Data are presented as the mean ± Standard derivatives (SD). Statistical significance was evaluated by Welch’s *t*-test.

Principal component analysis (PCA) of RNA-seq data from Caco-2 and MHCO cells cultured using iHACS revealed substantial differences and a clear separation of transcriptional patterns between both samples (Fig. 1B). The transcriptomes of Caco-2 and MHCO cells differed significantly, and among genes with an expression difference of > 1.5 fold change (adjusted *P*-value [*P*adj] < 0.05), we identified 4,942 and 4,824 genes that were enriched explicitly in Caco-2 and MHCO cells, respectively (Fig.1C). Furthermore, we visualized two non-overlapping classes of cell type-specific transcripts using hierarchical clustering, Caco-2 specific and MHCO cell signatures that were expressed in a mutually exclusive manner (Fig. 1D). Among the top 2,000 differentially expressed genes (DEGs), we observed several marker genes of functionally differentiated cells constituting the colonic epithelium: *Keratin 19 (KRT19*, colonocytes), *Mucin 2* (*MUC2*), *Trefoil factor 3* (*TFF3*, goblet cells), and *Chromogranin A* (*CHGA*, enteroendocrine cells) in the MHCO cell RNA-seq data (Fig. 1E, D). The stem cell marker gene, *Leucine rich repeat containing G protein-coupled receptor 5* (*LGR5*) was highly expressed in the cancer-derived cell line, whereas other markers of differentiated cells constituting the colonic epithelial tissue were highly expressed in MHCO cells (Fig. 1E). This finding was further supported by data showing that several known marker genes of the small intestine, such as *Alkaline Phosphatase, intestinal* (*ALPI*)*, Alanyl Aminopeptidase, membrane* (*ANPEP*), and *Apolipoprotein B* (*APOB*), were more highly expressed in Caco-2 than they were in MHCO cells (Fig. 1D, E).

Next, the DEGs that differed significantly between Caco-2 and MHCO cells (Fig 1D) were examined using gene ontology (GO) enrichment analysis. GO analysis of the genes upregulated in cultured Caco-2 cells showed enrichment of GO terms associated with the Wnt signaling pathway (GO:0016055) and different transport processes, including lipids (GO:0006869), organic anions (GO:0015711) and vitamins (GO:0051180, Fig. 2A). The GO enrichment analysis of Caco-2 cells showed enrichment of terms related to Wnt signaling pathway and lipids, organic anions, and vitamin transport processes. These findings are consistent with those previously reported on the identity of Caco-2 cells derived from a patient with colorectal cancer who had a mutation in the *APC* gene. *APC* is a negative regulator of Wnt signaling[22] that induces the expression of the target genes, *LDL Receptor Related Protein 1* (*LRP1)*[23], *Dickkopf-1* (*DKK1*)[24], and *Naked cuticle 1* (*NKD1*)[25]. Additionally, GO analysis revealed a significant association between the transporters of lipids (apolipoprotein family genes), organic anions (*D-allose ABC transporter membrane subunit* [*ASLC*]*/solute carrier organic anion transporter family member* [*SLCO*] family genes), and vitamins (*solute family carrier* [*SLC*] *family*, *NPC1 like intracellular cholesterol transporter 1* [*NPC1L1*], *ATP binding cassette subfamily G member 2* [*ABCG2*]) because Caco-2 cells are an extensively used *in vitro* assay for nutrient abruption studies[15] (Fig. 2B).

**Figure 2.**
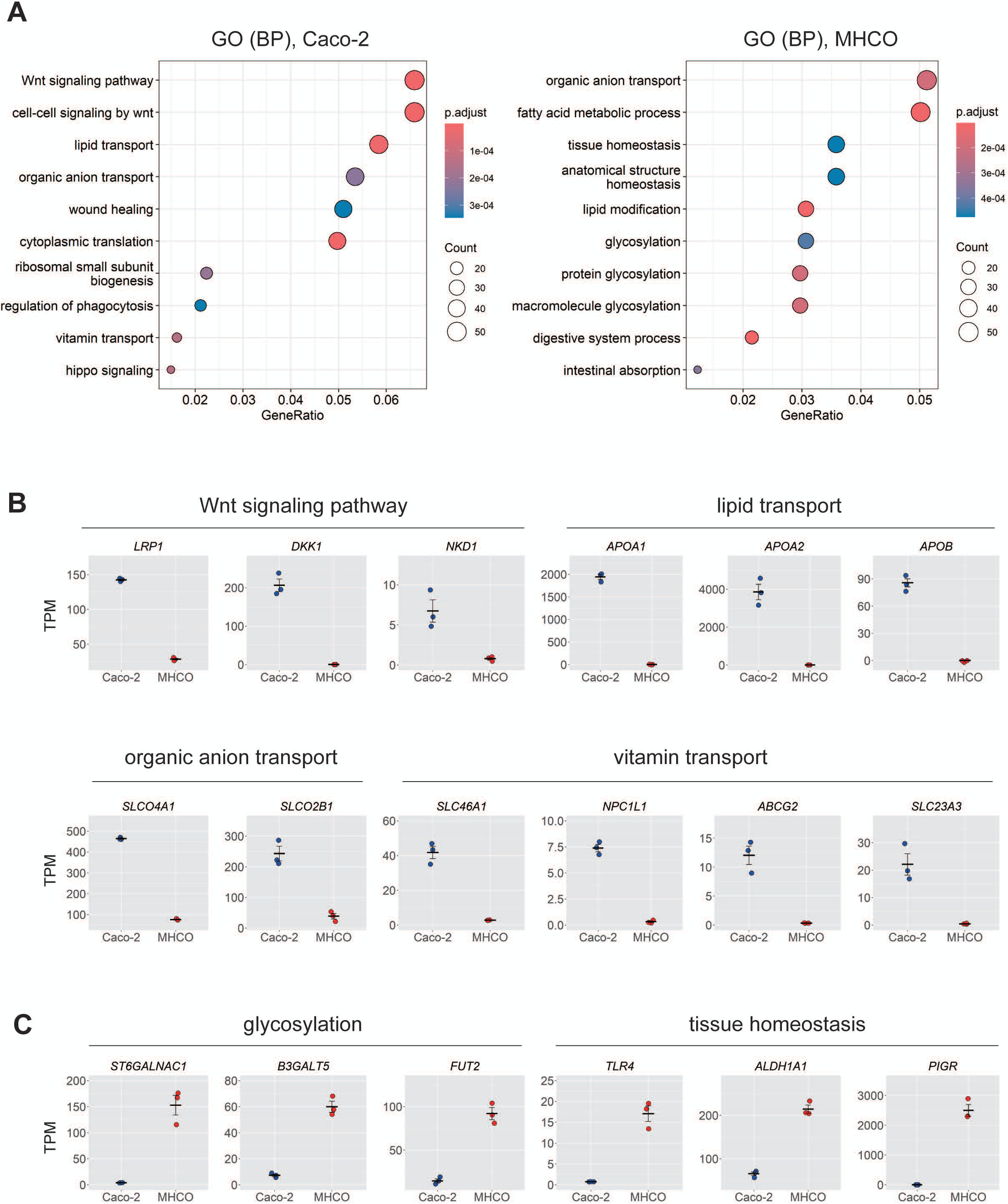
GO enrichment analysis of DEGs of the transcriptomes of MHCO and Caco-2. **(A)** Dot plot showing GO terms associated with DEGs for Caco-2 (left panel) and MHCO (right panel) identified in Figure 1D. **(B)** TPM values of representative genes for each GO term. Data are presented as the mean ± Standard derivatives (SD). Statistical significance was evaluated by Welch’s *t*-test.

In contrast, genes upregulated in MHCO cells showed a pronounced enrichment of GO terms related to tissue homeostasis (GO:0001894) involved in goblet cell marker genes (*MUC2* and *TFF3*) and different glycosylation processes, such as protein glycosylation (GO:0006486) and macromolecular glycosylation (GO:0043413) involved in the glycosylation of mucin proteins (*ST6 N-acetylgalactosaminide alpha-2,6-sialyltransferase 1* [*ST6GALNAC1*]*, beta-1,3-galactosyltransferase 5* [*B3GALT5*], and *Fucosyltransferase 2* [*FUT2*]; Fig. 1E and 2A and C).

Furthermore, GO analysis indicated the enrichment of tissue homeostasis genes (GO:0001894) exclusive to MHCO cells and involved in the mucosal immune system (Fig. 2A). Immune responses to gut bacteria, Toll-like receptor 4 (*TLR4*)[26], *Aldehyde dehydrogenase 1A1* (*ALDH1A1*)[27], and *Polymeric immunoglobulin receptor* (*PIGR*)[28], were detected in MHCO cells (Fig.2C). Collectively, the presence of these genes and enrichment of relevant GO confirmed that MHCO cells are more pertinent to observe the physiological response of the colonic epithelium than Caco-2 cells when those cells were cultured under semi-anaerobic conditions.

### Response of Caco-2 and MHCO cells to live anaerobes in the colon

Caco-2 cells have been used to analyze interactions between host cells and gut bacteria[29], and, therefore, we determined whether the responses of those cells to gut bacteria differed from those of MHCO cells. To test the physiological effects of commensal anaerobes in the colon, we cocultured Caco-2 or MHCO cells with a probiotic strain of *B. longum* using iHACS. The incubation time was recently extended to allow the coculture of the intestinal epithelium with anaerobic microbes[10]. Because *Bifidobacterium* is generally localized in the colon, we used healthy MHCOs in this study[30].

We confirmed that *Bifidobacterium longum* expression increased similarly in Caco-2 and MHCO cells cultured with iHACS (Fig. 3A, B). After 24 h, an inoculum of 1 × 10^7^ colony forming units (CFU)/mL of *B. longum* increased to a median of 6.98 × 10^8^ and 8.93 × 10^8^ CFU/mL in coculture with MHCO and Caco-2 cells, respectively (*P* = 0.26, Fig. 3B). RNA was extracted from Caco-2 and MHCO cells to assess their general responses to the probiotic bacteria. In the PCA of the characterized compound classes in Caco-2 and MHCO cells in the presence or absence of *B. longum* exposure (Fig. 3C), the first two principal components explained 33.8% (PC1) and 17% (PC2) of the total variation. PC1 indicated that MHCO cells were more distant from the clusters of Caco-2 cells, whereas PC2 showed the response of Caco-2 cells cocultured with *B. longum*. Samples of MHCO cells alone and cocultured with *B. longum* showed a low variation in each data set (Fig. 3C). Compared to the bacteria-free axenic culture, Caco-2 and MHCO cells incubated with *B. longum* showed distinct transcriptomic responses, as shown in the volcano plot of replicates exposed to identical conditions (Fig. 3D). After filtering for adj*P* > 0.05, the results showed that stimulation with *B. longum* affected the expression of 6,127 genes in Caco-2 cells by 1.5-fold, consisting of 3,062 and 3,065 upregulated and downregulated genes, respectively (Fig. 3D). Similarly, 2,886 genes were differentially expressed in MHCO cells, consisting of 1,407 and 1,479 upregulated and downregulated genes, respectively (Fig. 3D). Additionally, the overlap of upregulated DEGs between MHCO cells and Caco-2 was minimal, with only 269 commonly expressed genes (Fig 3E). Similarly, 296 DEGs exhibited decreased expression (Fig. 3E).

**Figure 3.**
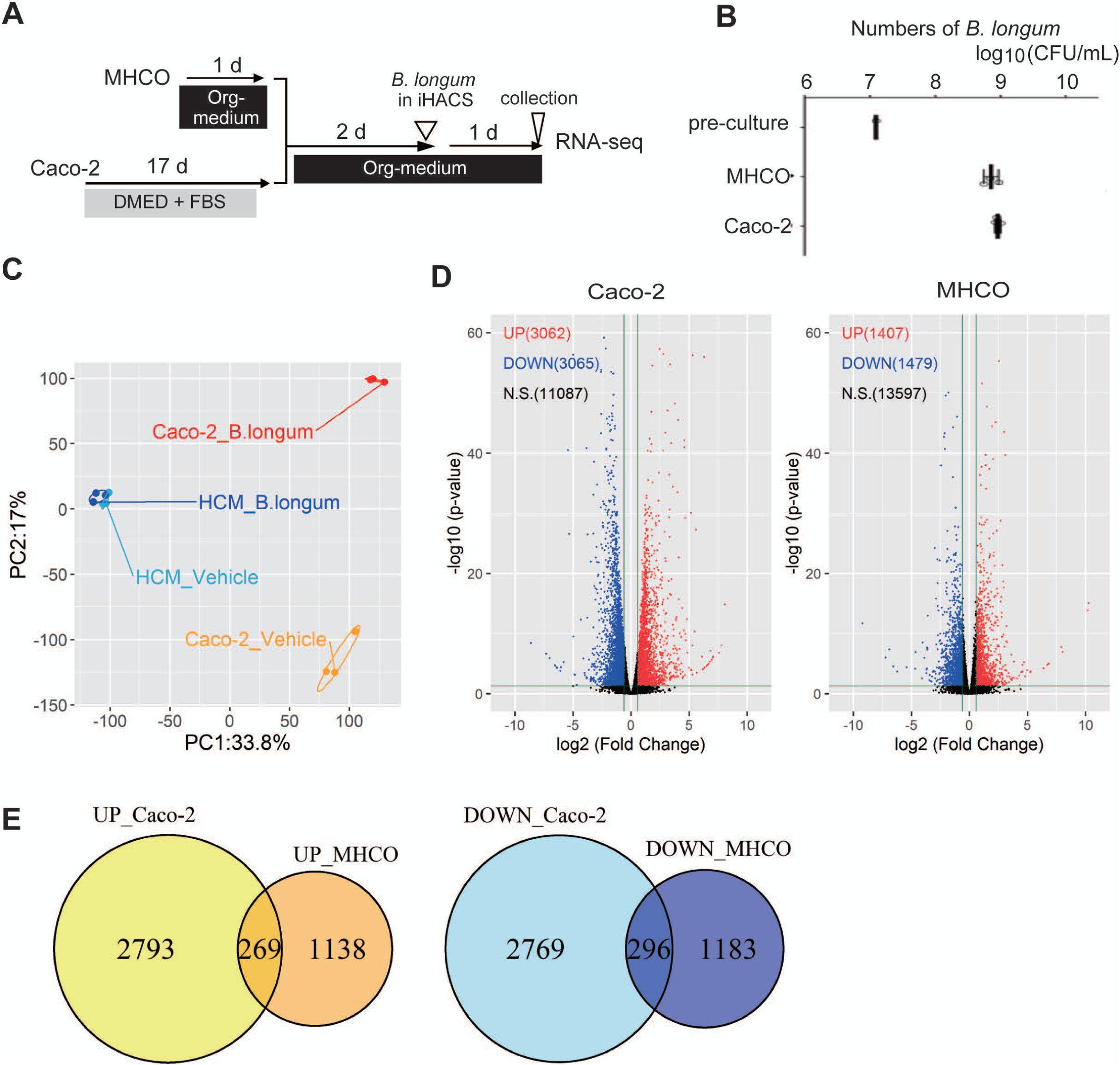
Differences in gene expression change among MHCO and Caco-2 after coculturing with *B. longum*. **(A-D)** A schematic representation of the experiment. **(B)** Colony-forming units (CFU) counts per milliliter of *B. longum* inoculated after 24-hour incubation with MHCO and Caco-2 using iHACS. **(C)** PCA of the transcriptomes of MHCO and Caco-2 cocultured with or without *B. longum*. Each plot represents a single transcriptome. Transcriptomes from three replicates were plotted. Ellipses indicate the 95% confidence interval. **(D)** Volcano plot depicting the upregulated and downregulated genes of Caco-2 (left panel) and MHCO (right panel) after coculturing with *B. longum*. The x- and the y-axis show the log_2_ transformed fold change and the -log_10_ transformed adjusted p-value. Red and blue dots indicate significantly upregulated and downregulated genes after coculturing with *B. longum* according to adjusted p-value <0.05 and log_2_ fold change >2. **(E)** Venn diagram comparing the significantly upregulated (left panel) and downregulated genes (right panel) in MHCO or Caco-2 after coculturing with *B. longum*.

The results of the GO analysis of genes that exhibited variations in expression in each cell are shown in Fig. 4. We identified GO terms derived from Caco-2 or MHCO cell-specific gene expression variations following stimulation with *B. longum*. The DEGs that showed increased expression exclusively in MHCO cells were predominantly enriched in terms related to lipid metabolism, with the most enriched GO term being lipid transport (GO:0006869) (Fig. 4A). Genes downregulated in MHCO cells were associated with cell proliferation pathways such as nuclear division (GO:0000280) and chromosome segregation (GO:0007059, Fig. 4B). In Caco-2 cells, the most enriched and least downregulated GO terms were cytoplasmic translation (GO:0002181) and the Wnt signaling pathway (GO:0016055), respectively (Fig. 4C, D). Conversely, most enriched terms in the commonly upregulated genes were associated with lipopolysaccharide (GO: 0032496) and response to tumor necrosis factor (GO:0034612 and GO:0071356), whereas downregulated genes were related to double-strand break repair (GO:0006302) (Fig. 4E, F).

**Figure 4.**
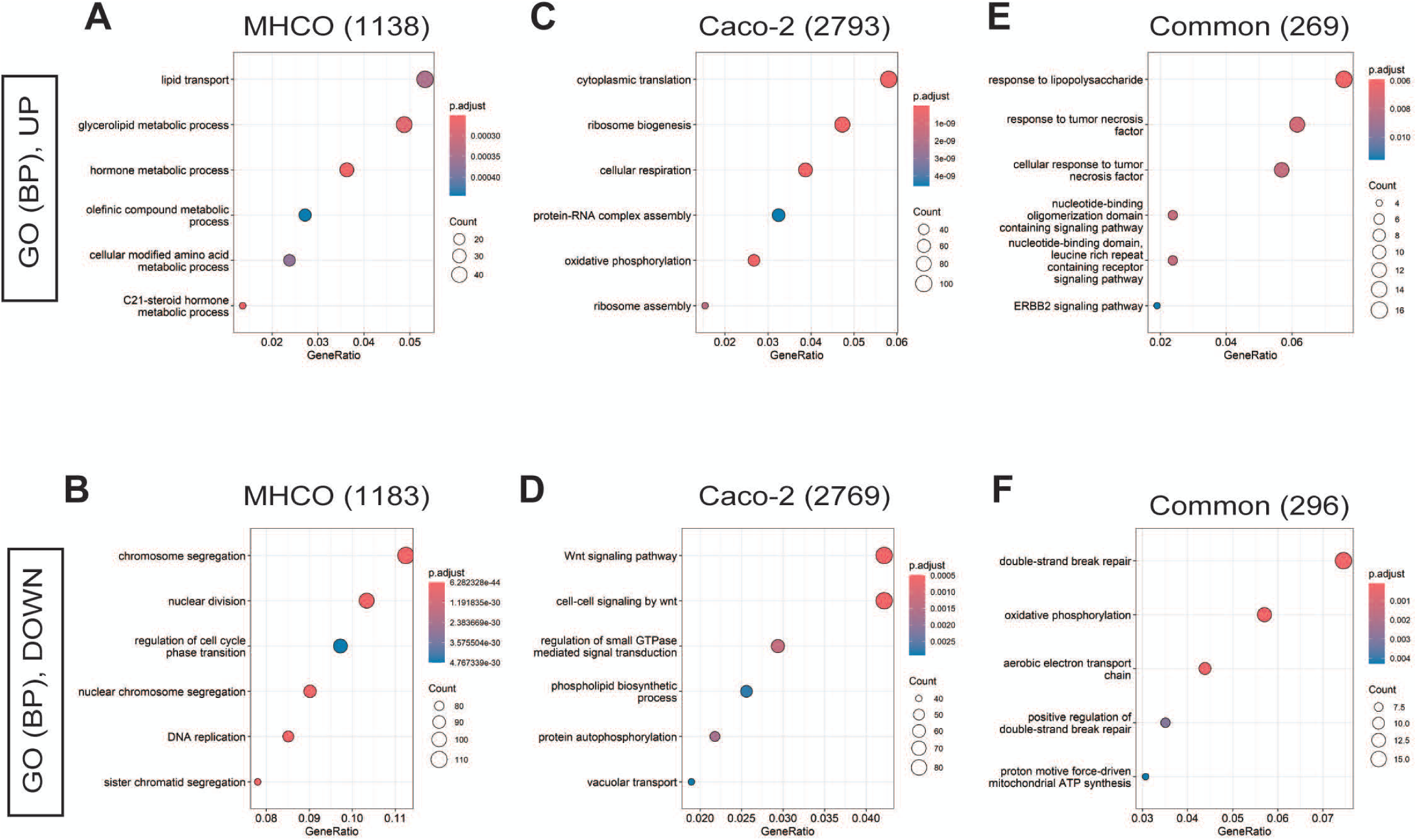
GO enrichment analysis of upregulated and downregulated genes in MHCO and Caco-2 after coculturing with *B. longum*. **(A-F)** Dot plot showing GO terms associated with upregulated (**A-C**) and downregulated genes (**D-F**) in MHCO (**A**, **D**), Caco-2 (**B**, **E**), and both (**C**, **F**) identified in Figure 3E.

### *B. longum* regulates the expression of transcription factors and genes involved in host metabolic pathway

Our analysis of the differentially induced gene expression in Caco-2 and MHCO cells following bacterial treatment showed a significant increase in the expression of genes involved in peroxisome proliferator-activated receptor (PPAR) signaling in MHCO cells (Fig. 5A). Furthermore, our transcriptome data showed that the expression of *PPARG* and *PPARA* were upregulated and downregulated in MHCO and Caco-2 cells, respectively following stimulation with *B. longum* (Fig. 5A, B). Genes of signaling pathways associated with PPARs, ketogenesis, lipid transport, cholesterol, fatty acid transporter/oxidation, adipocyte differentiation, and gluconeogenesis were upregulated in MHCO cells but not in Caco-2 cells (Fig. 5A).

**Figure 5.**
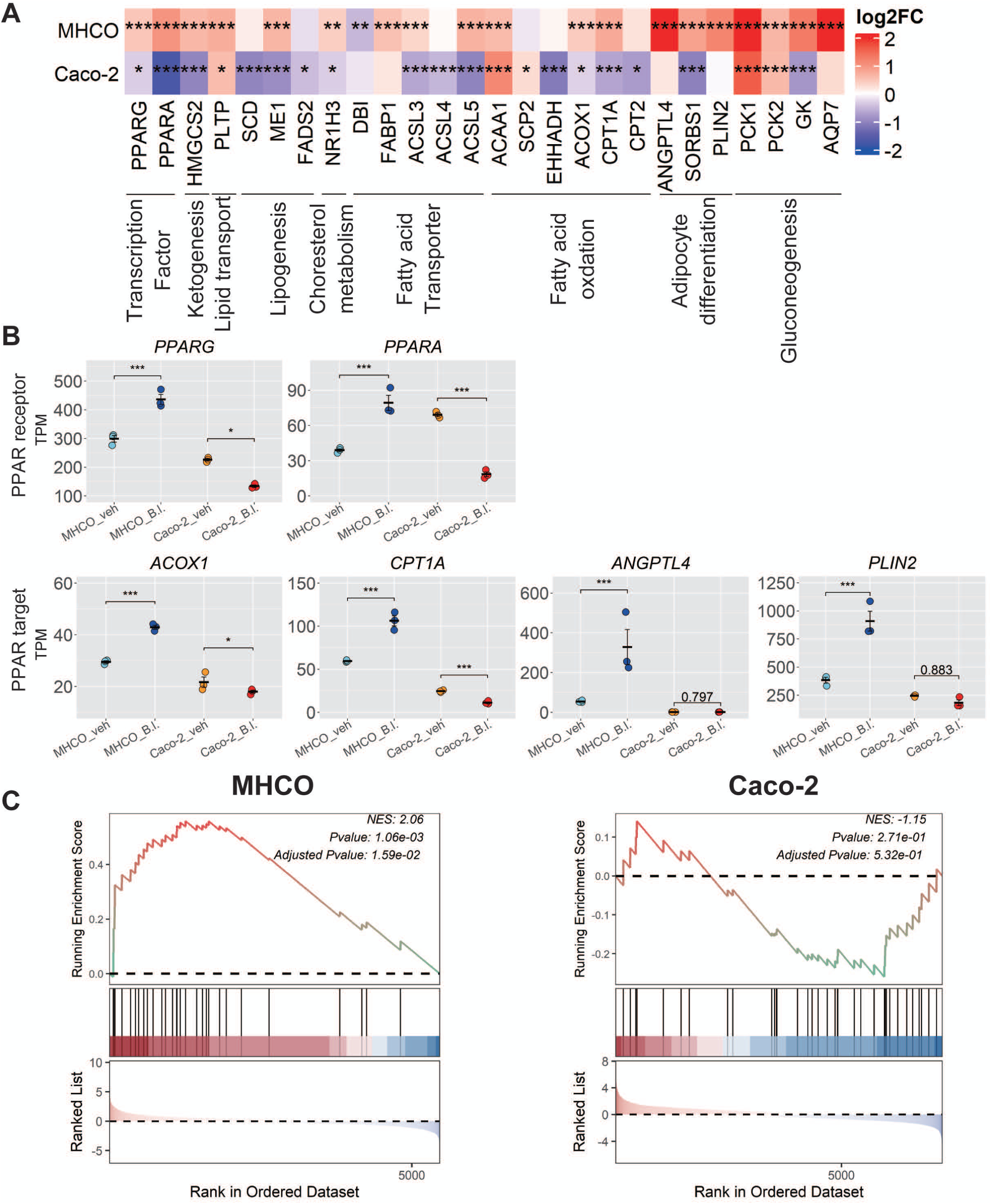
*B. longum* regulates the PPAR signaling pathway. **(A)** Heatmap of relative expression of PPAR signaling-related genes by co-cultivation of *B. longum* in Caco-2 and MHCO. Statistical significance was evaluated by a Deseq2 (*adjusted p-value <0.05, ** adjusted p-value <0.01, *** adjusted p-value <0.001). **(B)** TPM values of representative PPAR receptor and its target genes. Data are presented as the mean ± Standard derivatives (SD). Statistical significance was evaluated by Welch’s *t*-test. **(C)** Enrichment plot of genes in Ppar Signaling Pathway (KEGG) for MHCO (left panel) and Caco-2 (right panel). Each vertical bar represents a gene, and genes enriched in upregulated and downregulated after coculturing with *B. longum* are at the left and right part of the graph, respectively. The normalized enrichment score (NES), the p-value, and the false discovery rate (Adjusted p-value) were indicated in the insert.

To validate the upregulation of PPAR signaling in MHCO cells cocultured with *B. longum*, we focused on the expression of the PPARA and PPARG target genes. In MHCO cells coculturing with *B. longum*, *a fatty acid oxidation gene acyl-CoA oxidase 1* (*ACOX1*), a direct target gene of PPARA in the intestine[31], was elevated (Fig. 5B). *Angiopoietin like 4* (*ANGPTL4*), another identical target gene of PPARG in colonocytes[32], was also upregulated in MHCO cells (Fig. 5B). As expected, gene set enrichment analysis (GSEA) indicated that MHCO cells, but not Caco-2 cells cocultured with *B. longum* were enriched in genes related to PPAR signaling (normalized enrichment score [*NES*] = 2.06, *adj P* = 0.016 in MHCO cells, *NES* = -1.15, adj*P* = 0.53, Fig. 5C). These data indicated that live *B. longum* has a potential to activate PAPR signaling on normal physiological conditions of colonic epithelium.

### Differential inflammatory response to MHCO and Caco-2 cells to *B. longum* stimulation

Next, we characterized the reaction of each of the cell lines to microbial stimulation by determining their immune responses following stimulation and further investigated gene expression profiles in the production of cytokine and chemokine molecules and their regulators[33]. GSEA revealed that Caco-2 and MHCO cells stimulated by *B. longum* were commonly enriched for genes associated the NF-κB signaling pathway (NES = 1.92, P < 0.01, adj*P* < 0.05 in MHCO cells; NES = 2.43, *P* < 0.001, adj*P* < 0.001 in Caco-2 cells; Fig. 6A). To assess the responses of the Caco-2 and MHCO cells comparatively, we calculated the vehicle control and *B. longum*-induced log_2_ fold change (FC) for each gene. As expected, the responses induced by five chemokine genes (*C-C motif chemokine ligand 20* [*CCL20*], *C-X-C motif chemokine ligand 1* [*CXCL1*], *CXCL2*, *CXCL3*, and *CXCL8*) implicated in bacterial sensing[34] showed more pronounced FC values in Caco-2 cells than in MHCO cells (Fig. 6B, C).

**Figure 6.**
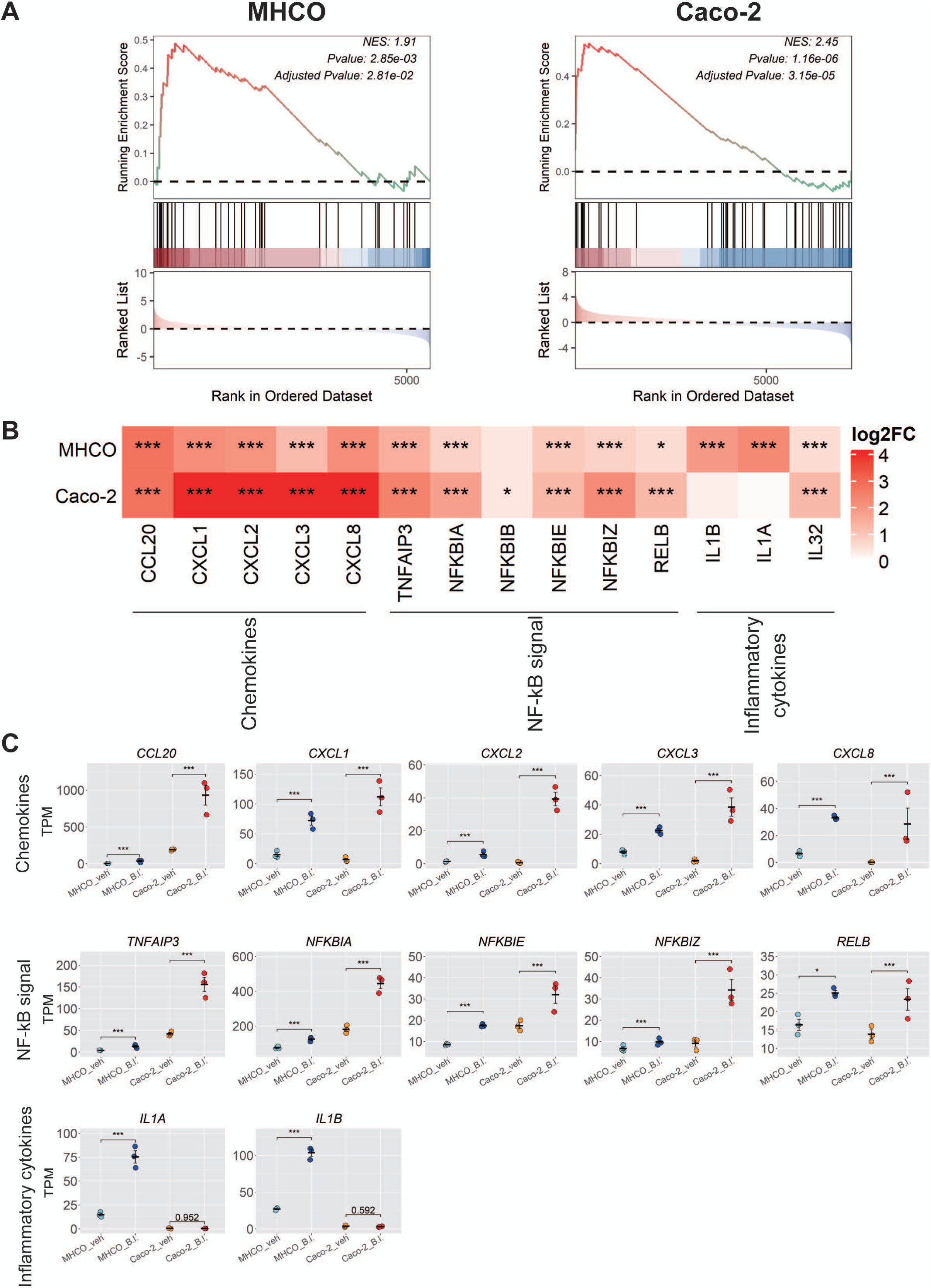
*B. longum* regulates inflammatory response differently in MHCO and Caco-2. **(A)** Enrichment plot of genes in Nf-Kappa B Signaling Pathway (KEGG) for MHCO (left panel) and Caco-2 (right panel). **(B)** Heatmap of relative expression of inflammatory response-related genes by co-cultivation of *B. longum* in Caco-2 and MHCO. Statistical significance was evaluated by a Deseq2 (*adjusted p-value <0.05, ** adjusted p-value <0.01, *** adjusted p-value <0.001). **(C)** TPM values of representative chemokine, NF-κB related, and inflammatory cytokine genes. Data are presented as the mean ± Standard derivatives (SD). Statistical significance was evaluated by Welch’s *t*-test.

Moreover, bacterial inoculation of Caco-2 and MHCO cells enhanced the expression of NFKB signaling target genes (*TNFAIP3*, *NFKB inhibitor alpha* [*NFKBIA*], NFKB inhibitor epsilon [*NFKBIE*], NFKB inhibitor zeta [*NFKBIZ*], RELB proto-oncogene, NF-κB subunit [*RELB*])[35] (Fig. 6B, C). Furthermore, *Interleukin 1A* (*IL-1A*) and *IL-1B*, inflammatory cytokine genes crucial for inflammation and infection defense, were significantly upregulated following coculturing of *B. longum* and MHCO cells specifically [36] (Fig. 6B, C). Those results suggest that a probiotic microbe, *B. longum,* raises the level of immune surveillance.

### *B. longum* upregulates intestinal barrier function in MHCO cells

We investigated the effect of Bifidobacteria on the intestinal barrier function, by examining the effects on expression patterns of relevant genes. Our RNA-seq results revealed substantial differences in expression patterns of genes related to the mucosal barrier function between Caco-2 and MHCO cells in response to *B. longum*. The tight junction-related genes, *Claudin 3* (*CLDN3*), *CLDN4,* and *Occludin* (*OCLDN*), were upregulated in both Caco-2 and MHCO cells following exposure to *B. longum*. In contrast, other related genes, *CLDN7* and *Tight junction protein 3* (*TJP3*), were only upregulated in MHCO cells, whereas *CLDN2* was upregulated in Caco-2 cells (Fig. 7A, B). Other components of the adherens junction (Cadherin 1 [*CDH1*], *CDH17*, *Epithelial cell adhesion molecule* [*EPCAM*]) or desmosomes (*Desmoglein 2* [*DSG2*] and *Dystonin* [*DST*]) were linked to a set of genes that were upregulated in *B. longum*-stimulated MHCO cells and conversely downregulated in Caco-2 cells (Fig. 7A, C, D). The polarity of intestinal epithelial cells plays a central role in establishing the barrier function for symbiotic relationships with microbiota[37]. The *B. longum*-induced expression pattern of *Integrin subunit α6* (*ITGA6*), a significant factor in the epithelium and basement membrane interaction, also differed between MHCO and Caco-2 cells (Fig. 7A, E).

**Figure 7.**
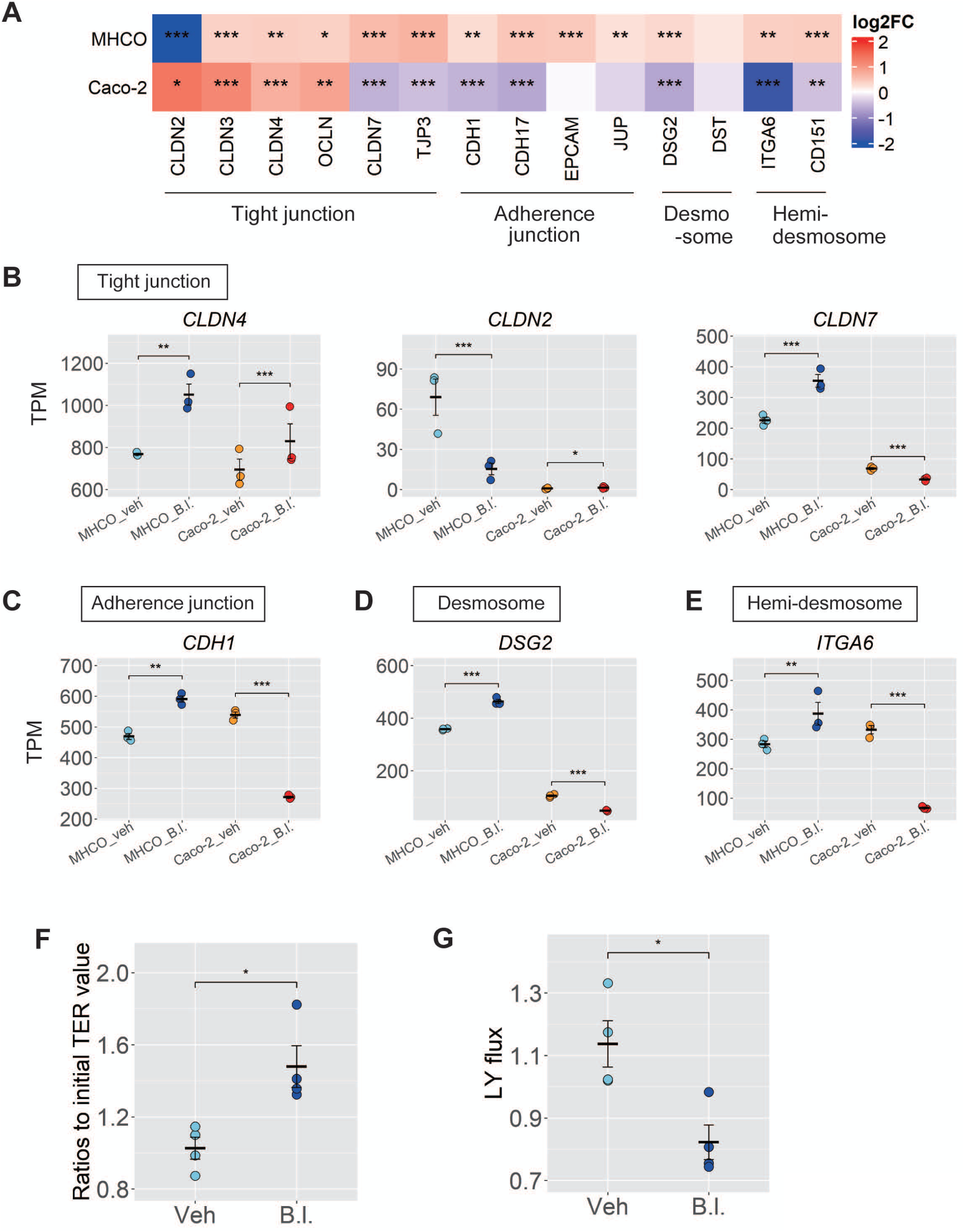
*B. longum* regulates intestinal barrier function. **(A)** Heatmap of relative expression of cell junction-related genes by co-cultivation of *B. longum* in Caco-2 and MHCO cells. Statistical significance was evaluated by a DESeq2 (*adjusted p-value <0.05, ** adjusted p-value <0.01, *** adjusted p-value <0.001). **(B)** TPM values of representative tight junction (upregulated in both MHCO and Caco-2 cells, specifically upregulated in MHCO), adherence junction, and hemidesmosome-related genes. Data are presented as the mean ± Standard derivatives (SD). Statistical significance was evaluated by Welch’s *t*-test. **(C, D)** Relative TEER value adjusted by the initial value (**C**) and permeability of Lucifer Yellow (LY) from apical to basolateral compartment (**D**) in MHCO cocultured with *B. longum* versus vehicle control. Data are expressed as means ± SD. Statistical significance was evaluated using Welch’s t-test.

To investigate whether increased gene expression induced by *B. longum* was associated with improved barrier function, we measured the transepithelial electrical resistance (TEER) in MHCO cells. Compared to the vehicle control, TEER was significantly increased in cells cocultured with *B. longum* (Fig. 7D). We also evaluated the paracellular permeability of the colonic epithelium using the Lucifer yellow (LY) flux assay with the fluorescent molecule LY, which does not interact with cell components[38]. Consistent with the changes in TEER, LY flux was downregulated when MHCO cells were cultured with *B. longum* (Fig. 7E). Overall, these results confirmed the previous studies in MHCO cells with live *B. longum* in hemi-anaerobic culture conditions that probiotic *Bifidobacteria* enhance intestinal barrier integrity through PPARG/STAT3 pathway.

## Discussion

In this study, we compared gene expression data between Caco-2 and MHCO cells cultured under hemi-anaerobic conditions to provide valuable datasets for investigating host cells and anaerobic bacteria in the colon. We demonstrated the presence of all cell types that form the colonic epithelium, including stem, goblet, and enteroendocrine cells and colonocytes among the MHCO cells cultured under hemi-anaerobic conditions. In contrast, the gene expression pattern of Caco-2 cells was similar to that of epithelial cells of the small intestine, as shown previously[39]. Therefore, MHCO cells may be a physiologically appropriate model for studying probiotic activities, including of *Bifidobacterium* and *Akkermansia*, in the colon[30, 40].

The Caco-2 cell line generated from a patient with colorectal cancer is the most commonly used in studying the physiological functions of the human intestinal epithelium. These cells spontaneously transform to differentiate and form monolayers of polarized cells with functions similar to those of enterocytes in the small intestine when grown under confluent conditions in Boyden chambers. Consequently, Caco-2 cells have been widely used to investigate the metabolism and absorption of drugs and nutrients[15, 41]. The intestinal environment where enterobacteria are localized is physiologically hypoxic, which affects gene expression patterns through the chromatin state and metabolic pathways[42].

Gnotobiotic mice have been the primary animal model used in previous studies of the effect of microbes on hosts. Although studies using in vivo mouse models provide important insights into the physiology of host-microbiome interactions, they are unable to account for significant differences between mice and humans because bacteria exhibit strong species specificity[43]. Conventionally, cell lines derived from patients with cancer have been used to study microbial responses. In this study, healthy primary colonic epithelial cells cultured as organoids and a conventional colorectal cancer-derived cell line were used to examine the physiological response of the host epithelium to microbiota. The gene expression patterns in the DEG analysis in our study indicated that healthy human colonic epithelium derived from organoids was more robust against gut bacteria than the cancer-derived Caco-2 cell line was. These results demonstrated the observed low incidence of overlapping upregulated DEGs between MHCO and Caco-2 cells and decreased expression of DEGs, indicating that the impact of coculturing with *B. longum* on gene expression was not conserved across these two cell types.

Caco-2 cells have been used to assess the efficacy of probiotics and intestinal bacteria in evaluating mucosal barriers and inflammatory responses. However, two independent reports demonstrated the immunoreactivity of *Faecalibacterium prausnitzii* (recently reclassified as *Faecalibacterium duncaniae*) in host cells using Caco-2 cells or primary colonic epithelium (organoid)[12, 44]. To the best of our knowledge, this study is the first to evaluate the effect of gut bacteria on Caco-2 cells and colonic organoids incubated under matched culture and hemi-anaerobic conditions.

In this study, the expression of various inflammation-related genes was similarly induced by *B. longum* in Caco-2 and organoid-derived healthy epithelial cells. However, the fold-change ratio was higher in Caco-2 cells than it was in the organoid cells. This probably occurred because Caco-2 cells lack goblet cells and poorly express genes such as *MUC2*, which renders the barrier function of the intestinal epithelium vulnerable. Moreover, the bacteria may have more readily caused inflammation in the Caco-2 cells than in MHCO cells. Therefore, we focused on immune responses to characterize the reaction of Caco-2 and MHCO cells to microbial stimulation because genes commonly upregulated in these cell lines show pronounced enrichment of GO terms associated with inflammation. The intestinal epithelium interfaces directly with the external environment and acts as a constant frontline defense for the immune response[45]. Epithelial cells play an essential role in preventing bacterial invasion by producing cytokines and chemokines that determine the nature of the immune response by controlling immune cells. These facts prompted our investigation of gene expression profiles in the production of immune-related molecules and their regulators, including NFKB inhibitors and TNFAIP[33]. The results suggest that the probiotic microbe, *B. longum,* raised levels of immune surveillance.

In addition, our GSEA results showing the enrichment of genes related to PPAR signaling in bacterial-stimulated MHCO but not Caco-2 cells led us to conclude that live *B. longum* has the potential to activate PAPR signaling under normal physiological conditions in the colonic epithelium. These observations further confirmed that MHCO cells are more pertinent for observing the physiological response of the colonic epithelium than Caco-2 cells are when cultured under semi-anaerobic conditions.

Moreover, the results of the TEER analysis corroborated the relationship between *B. longum*-induced increase in gene expression and improved barrier function in MHCO cells. These observations further confirmed the findings of previous studies in MHCO cells cocultured with live *B. longum* under hemi-anaerobic culture conditions, which showed that probiotic *Bifidobacteria* enhanced the intestinal barrier integrity through the PPARG/signal transducer and activator of transcription 3 (STAT3) pathway. Previous animal and clinical studies have revealed that *Bifidobacterium*, a probiotic, positively regulates the barrier function of mucosal epithelial cells[46].

Epithelial cells are held together by strong anchoring junctions that include tight junctions, adherens junctions, and desmosomes, which are formed by transmembrane adhesion proteins[47]. The expression of *CLDN4*, a component of tight junctions, has been shown to be upregulated in the colonic epithelial cells of mice administered *Bifidobacteria*[48]. Similarly, our investigation using human colonic organoids cocultured with *B. longum* also showed the upregulation of *CLDN4*. Contrastingly, several molecules related to epithelial cell junctions, such as *CLDN2*, showed conflicting expression patterns in Caco-2 and MHCO cells stimulated with *B. longum*. *CLDN2* is a pore-forming CLDN that permeates water and ions and has been reported to be elevated in patients with inflammatory bowel disease or neonatal necrotizing enterocolitis[49–52]. Furthermore, we found that *CLDN7*, a tight junction-related gene, was upregulated only in MHCO monolayer cells cocultured with *B. longum*. Increased *CLDN2* and decreased *CLDN7* levels have previously been observed in the gastrointestinal tracts of patients with ulcerative colitis[53], and *CLDN7* knockout mice develop colitis spontaneously[54]. In this study, to the best of our knowledge, we provide the first proof that *Bifidobacterium* modulates *CLDN*7 expression in the physiologically normal human colonic epithelium.

The polarity of intestinal epithelial cells plays a central role in establishing a barrier function for symbiotic relationships with microbiota[37]. Hemidesmosomes, which are composed of ITGA6T and are a significant factor in the interaction between the epithelium and basement membrane, are expressed at specific junctions in the basal portion of intestinal epithelial cells[55]. In addition to other factors, improvement of the mucus barrier function is a known molecular mechanism in coculture systems. A novel finding of this study is that more genes were altered by bacterial stimulation in Caco-2 cells than they were in the organoids. While both organoids and Caco-2 cells share certain variable gene sets, each cell line exhibited a unique variation in the expression of genes linked to inflammation and the mucosal barrier, which are well-known roles of *Bifidobacterium*. However, the potential usefulness and applicability of *Bifidobacterium* needs to be further investigated using appropriate *in vitro* culture systems.

## Methods

### Colonic organoid culture

Clinical samples for healthy human colonic organoid establishment and coculturing microbes were obtained from patients at Gunma University Hospital with informed consent after study approval by the ethical committees (HS2022-054). The human colonic organoid was cultured as in the previous report with minor modifications[10]. Briefly, three-dimensional colonic organoids were maintained with Modified human colonic organoid (MHCO) medium, consisting of advanced Dulbecco’s modified Eagle’s medium (DMEM)/F12 supplemented with penicillin/streptomycin, HEPES, Glutamax, B-27 Supplement (Thermo Fisher Scientific, Waltham, MA), 1 mM N-acetylcysteine (Sigma-Aldrich), 50 ng/mL recombinant mouse epidermal growth factor (Thermo Fisher Scientific), 100 ng/mL mouse recombinant noggin (Peprotech), 1μg/mL human recombinant R-spondin1 (R&D), 100 ng/mL recombinant human insulin-like growth factor-1 (BioLegend, San Diego, CA), 50 ng/mL recombinant human fibroblast growth factor-basic (FGF-2) (PeproTech) and conditioned medium containing Wnt3A. The human colonic organoid was derived from a nonpathological biopsy and confirmed by genomic sequencing analysis that it has no driver mutations related to colorectal cancer, as shown in the previous paper^2^. Organoids were passed approximately every 5-7 days by physical dissociation using fire-polished Pasteur pipettes. To generate MHCO cells, ThinCert culture inserts (24-well insert, 0.4 μm pore polyester membrane; Greiner bio-one, Kremsmunster, Austria) were coated with 4% Matrigel diluted with advanced DMEM/F12 medium and incubated at 25°C for 30 minutes, then Matrigel solution was removed. The membrane was dried in a tissue-culture hood for 15 minutes. Human colonic organoids were cultured for 3 to 5 days before being used to plate into monolayer culture in MHCO medium. Three-dimensional cultured organoids were treated with TrypLE Express (Thermo Fisher Scientific) to dissociate into single cells. The cells were resuspended to 1× 10^6^ cells/mL in MHCO medium containing 10 μM Y-27632 (FUJIFILM Wako Pure Chemical Corporation, Osaka, Japan), and 200 uL of cell suspension was added into the transwell inserts. After 1 day of monolayer culture, MHCO without Y-27632 was used, and the medium was changed every 2 days. When MHCO cells were cultured in the inserts, MHCO without penicillin/streptomycin was used for their maintenance.

### Caco-2 cell culture

The cell line Caco-2 maintained in our group was confirmed by the Cell Line Authentication Service (ATCC). Caco-2 cells were maintained with the medium consisting of DMEM (high glucose) with L-Glutamine and Phenol Red (FUJIFILM Wako Pure Chemical Corporation) supplemented with 10% heat-treated FBS (Hyclone, Logan, UT), MEM Non-Essential Amino Acids Solution (Nacalai tescue, Kyoto, Japan), and penicillin/streptomycin. Caco-2 cells were passaged approximately 3-5 days before reaching 80% confluency using 0.25% Trypsin-EDTA (Gibco, Waltham, MA, USA) and seeded at 2-4 x10^4^ cells/cm^2^. 10^5^ cells were seeded in transwell culture insert coated Matrigel solution described above. Cells were cultured with the 10% FBS-containing DMEM medium for 17 days. Before the coculture experiment, the culture medium was changed to MHCO medium without antibiotics and cultured for 2 days.

### Bacterial culture

*B. longum* subsp. *longum* JCM1217^T^ (*B. longum*) was obtained from the Riken BioResource Research Center (JCM, Tokyo, Japan). This strain was cultured in modified GAM broth (Nissui Pharmaceutical, Gifu, Japan) for 4 hours at 37°C under anaerobic conditions using a BACTRON300 anaerobic chamber (Sheldon Manufacturing, Inc., Cornelius, USA). The culture medium was centrifuged (8,000 x g, 1 min, 25°C) and suspended to approximately 5.0 × 10^7^ CFU/mL in a deoxygenated WENRAIF medium supplemented with 100mM HEPES in an anaerobic chamber. Then, 200 uL bacterial suspension was added at the apical side of the transwell in an anaerobic chamber and capped in butyl gum caps. Then, the basolateral side’s medium was cultured in a CO2 incubator for 24 hours. The number of colony-forming units (CFU) on TOS propionate agar (Yakult Pharmaceutical Industry, Tokyo, Japan) was used to calculate the number of live *B. longum*.

### RNA isolation and sequencing

Total RNA was extracted using the RNeasy Mini Kit (Qiagen) with an RNase-Free DNase Set (Qiagen). Library preparation was performed using Illumina Stranded mRNA Prep and IDT® for Illumina® RNA UD Indexes Set A-B Ligation (Illumina, San Diego, CA, United States). The concentration and quality of the extracted RNA and adapter-tagged sequence library were calculated using Agilent RNA 6000 Nano and Agilent High Sensitivity DNA Kits (Agilent Technologies, Santa Clara, CA, USA), respectively. Sequences were obtained using the NextSeq 1,000 system with the Illumina NextSeq 1000/2000 P2 Reagent kit (100 cycles) (Illumina). Read files were trimmed and mapped to the human reference genome (GRCh38.p.13) using Trim Galore! (ver.0.6.4, https://github.com/FelixKrueger/TrimGalore) and HISAT2 (ver.2.2.1) (Graph-Based Genome Alignment and Genotyping with HISAT2 and HISAT-genotype - PMC (nih.gov)) with default settings[56]. Transcript assembly, GTF document merging, and transcript quantification were performed using Stringtie (ver.2.2.0) (StringTie enables improved reconstruction of a transcriptome from RNA-seq reads - PubMed (nih.gov))[57]. A matrix of read counts mapped to genes was extracted using prepDE.py (http://ccb.jhu.edu/software/stringtie/dl/prepDE.py3).

### Data analysis and visualization

Normalization from read counts and identifying differentially expressed genes (DEGs) were performed via iterative differential expression analysis (DESeq2) by iDEP1.1 software. Adjusted p-value <0.05 and log_2_ fold change >1.5 were defined as DEGs. Statistical analysis and data visualization were performed using R (ver.4.3.2). The principal component analysis (PCA) was performed using the ‘prcomp’ function with TPM matrix in the stats package (ver.3.6.2, https://www.rdocumentation.org/packages/stats/versions/3.6.2) with scaling. The hierarchical clustering and plotting of the heatmap were performed by the ‘Heatmap’ function in the ComplexHeatmap package (ver.2.18.0, https://github.com/jokergoo/ComplexHeatmap). Functional enrichment analysis of DEGs was performed using the ‘enrichGO’ function in the clusterProfiler package (ver.4.10.0, https://github.com/YuLab-SMU/clusterProfiler) and GO terms with p-value <0.05 and adjusted p-value <0.05 were considered significant. KEGG pathway enrichment analysis was performed using the ‘gseKEGG’ function in the clusterProfiler package, and the significantly enriched pathways were defined by nominal adjusted p-value <0.05.

### Measurement of the integrity of the cell monolayer

The integrity of the cell monolayer was assessed with relative TEER value and LY assay. TEER was measured by Millicell ERS-2 Voltohmmeter (Merck Millipore, USA) following the manufacturer’s instruction. Relative TEER was calculated by TEER value after Hemi-Anerobic cultivation for 24 hours relative to their initial value (0 hours). HBSS supplemented with 10mM HEPES, D-PBS(+) Preparation Reagent (Ca, Mg Solution) (Nacalai tesque, Kyoto, Japan) were used as transport buffers. After washing the monolayer with transport buffer, 300uL of Lucifer yellow solution (300uM prepared in transport buffer) and 1000uL of transport buffer were added to the apical and basolateral sides of the monolayers, respectively. The monolayers were incubated for 60 minutes in a CO2 incubator. After incubation, 200uL of apical and basolateral solution was collected in the 96-well black assay microplate, and the fluorescence was measured at an emission wavelength of 538 nm and excitation at 485 nm using Perkin Elmer Enspire 2300 Multi-mode Microplate Reader with EnSpire Workstation (version 4.13.3005.1482) n=4 biological replicates.

## Acknowledgment

The authors thank the members of Gastroenterological Surgery, School of Medical Faculty of Medicine at Gunma University for human sample collection and all members of the laboratory for Mucosal Ecosystem Design for their assistance in experiments and discussion. A Wnt3a-producing cell line was a kind gift from Hans Clevers (Hubrecht Institute).

## Funding

This work was supported in part by the Japan Agency for Medical Research and Development (AMED) grant (JP 23ae0121046), JST FOREST program (JPMJFR2161), Grants-in-Aid for the Japanese Society for the Promotion of Science (JSPS) KAKENHI (19H03455, 23H02713), a Grant-in-Aid for Challenging Research (Pioneering, 23K17415), the Astellas Foundation for Research on Metabolic Disorders, and the LOTTE Foundation.

## Author contributions

A.S., A.I., F.S., and T. Odamaki performed all the empirical experiments and analyzed the data. A. S., A. I., K. Y., E. M., and T. Oda. performed and analyzed the RNA-seq data. S.U., T. Okada, T.Y., and H.S. provided tissue specimens. N.S. conceived and conducted the project. A.S. and N.S. wrote the manuscript. All authors discussed the results and edited the manuscript.

## Conflict of interest

A. S. and T. O. are employees of Morinaga Milk Industry Co. Ltd. The remaining authors disclose no conflicts.

## Notes

### Summary of Updates

Changed the description of material method

